# Heat stress drives opposing redox shifts in temperate versus tropical *Drosophila melanogaster* embryos

**DOI:** 10.64898/2026.06.30.733001

**Authors:** Thomas S. O’Leary, Brent L. Lockwood

## Abstract

Redox balance is central to aerobic metabolism, yet acute heat stress can destabilize this balance by increasing metabolic rates and shifting the balance of critical electron carriers such as NADH. In early *Drosophila melanogaster* embryos, maintaining redox balance is particularly critical as embryos undergo a developmental redox shift and rely on oxidative phosphorylation to power nuclear divisions. Here, we assayed six isofemale *D. melanogaster* lines from temperate (Vermont, USA; France; Japan) and tropical (St. Kitts; Ghana; India) climates to assess metabolic responses to heat in heat-sensitive versus heat-tolerant embryos. We used untargeted LC–MS to measure 33 metabolites and the major redox couples (NADH/NAD^+^, NADPH/NADP^+^, and GSH/GSSG) at 25°C and after a 32°C heat shock. In all embryos, heat shock induced shared shifts in metabolic profiles, with increases in nucleotide monophosphates (*e.g.*, AMP, CMP, and GMP) and amino acids (*e.g.*, alanine, glutamic acid, serine). In contrast, redox metabolites diverged by region: heat-sensitive temperate embryos shifted toward a more oxidized state (46.6% decrease in NADH/NAD^+^ ratio and 4-fold increase in oxidized glutathione), while heat-tolerant tropical embryos maintained glutathione balance and increased the NADH/NAD^+^ ratio by 52.9%, indicating a more reduced state. These patterns are consistent with higher NADH oxidation and greater oxidative stress (inferred from oxidized glutathione) in the temperate embryos, versus better maintenance of redox balance in tropical embryos. Together, our results suggest that maintaining redox balance is a key determinant of acute heat tolerance, and healthy development overall, during early embryogenesis.

## Introduction

Many of the biochemical reactions that power life involve the transfer of electrons through reduction-oxidation (redox) reactions (Marques, 2024). This includes oxidative phosphorylation, which utilizes electron carriers such as NADH and FADH_2_ to fuel the electron transport system (ETS) and generate ATP (Somero et al., 2017). Reactive oxygen species (ROS), highly reactive oxygen-containing molecules with important cellular signaling functions, are a normal byproduct of this process (Buetler et al., 2004; Liu et al., 2002; Murphy, 2009; Sies and Jones, 2020; Syal et al., 2020; Zhang and Wong, 2021). But acute heat stress can create excessive ROS production (Belhadj Slimen et al., 2014; Han et al., 2023; Jimenez et al., 2016) which can damage DNA, lipids, proteins, and otherwise disrupt cellular functioning (Ezraty et al., 2017; Sharma et al., 2012; Sies and Cadenas, 1985). These negative effects are combatted by various antioxidant molecules, enzymes, and pathways (e.g., vitamin C, catalase, glutathione redox cycle) (Ferreira-Cravo et al., 2023; Halliwell, 2022; Jia et al., 2011; Lu and Holmgren, 2014; Vašková et al., 2023; Weydert and Cullen, 2010), yet severe heat stress can overwhelm antioxidant capacity and shift the cellular redox state (Rossi et al., 2024; Skjærven et al., 2013; Subedi et al., 2018).

Early embryos rely on the breakdown of maternally supplied glycogen and triglycerides to power ATP production (Tennessen et al., 2014) with oxidative phosphorylation reactivated following fertilization (Eller et al., 2025). Mitochondrial-supplied ATP is critical to supporting the rapid syncytial divisions that characterize early embryonic development, with even brief inhibition of the electron transport system disrupting key events during nuclear division (Chowdhary et al., 2017). Although the link between mitochondrial function and upper thermal limits is debated (Chung and Schulte, 2020), a rapid heat-induced imbalance in cellular redox state could plausibly impact mitochondrial output and disrupt embryogenesis (Murphy, 2009; Petrova et al., 2018).

Embryogenesis in a wide array of taxa (*e.g.,* urchins, frogs, zebrafish, and mice) involves tightly controlled changes in redox state (Dumollard et al., 2007; Han et al., 2018; Shapiro, 1991; Timme-Laragy et al., 2013). Shifts in redox balance are important in early *Drosophila melanogaster* development, where the oocyte-to-embryo transition is marked by changes in cellular redox state and proteome-wide changes in reactive cysteine residues driven by thioredoxins, leading to disruptions in meiosis, fertilization, and nuclear divisions in the early embryo (Petrova et al., 2018). This dependency on redox state may be particularly consequential in thermally variable contexts. Indeed, early *D. melanogaster* embryos are heat-sensitive relative to older life stages, unable to behaviorally thermoregulate, and operate near their upper thermal limits (Lockwood et al., 2018). Differences in heat tolerance between temperate and tropical embryos reflect thermal adaptation to climate (Lockwood et al., 2018; Nunez et al., 2026), with transcriptomic evidence further suggesting a potential adaptive role in maintaining redox homeostasis after heat stress (Mikucki et al., 2024). While the expression of oxidative stress genes is consistent with higher antioxidant capacity in heat-tolerant tropical embryos (Mikucki et al., 2024), the link between heat tolerance and cellular redox state in early *Drosophila* embryos is unknown.

Here, in temperate and tropical *D. melanogaster* embryos, we used untargeted liquid chromatography–mass spectrometry (LC–MS) to measure the heat-induced response of major redox couples (NAD(H), NADP(H), and GSH/GSSG) and other core metabolites. By measuring cellular redox state, we avoided the difficulty of measuring ROS directly (Kalyanaraman et al., 2012; Murphy et al., 2022), and we were able to determine the extent to which redox balance is correlated with heat tolerance. We tested whether (i) redox metabolite levels were predictive of acute embryonic heat tolerance, and (ii) whether there was a region-specific (i.e., temperate vs. tropical) shift in redox state following acute heat stress. Given the regional differences in heat tolerance and the potential association between heat stress and oxidative stress (Belhadj Slimen et al., 2014; Chung and Schulte, 2020; Mikucki et al., 2024), we predicted that heat-sensitive temperate embryos would show a unique shift towards a more oxidized state at sublethal temperatures.

## Results

### Higher acute heat tolerance in tropical embryos

We measured hatching success after a 45-minute acute heat shock in six *D. melanogaster* isofemale lines from temperate and tropical regions. Four of these lines (Vermont, USA; Chiapas, Mexico; Accra, Ghana; and Mumbai, India) were previously characterized by (Lockwood et al., 2018), and we added two new lines from Montpellier, France; and Shiojiri, Japan to extend temperate sampling beyond North America (**Fig. 1A**). As predicted from previous data, tropical embryos exhibited higher heat tolerance than temperate embryos, with a mean LT_50_ (the temperature at which 50% of embryos fail to hatch) of 36.0°C compared to 34.5°C for temperate lines (**Fig. 1B**). Embryonic LT_50_ was strongly predicted by the maximum temperature of the warmest month of each collection location (WorldClim bioclimatic variable 5): LT_50_ increased by 0.23°C for each 1°C rise in maximum temperature of the warmest month (slope = 0.227, S.E. = 0.020, R^2^ = 0.97, p = 0.0003; **Fig. 1C**).

**Figure 1.**
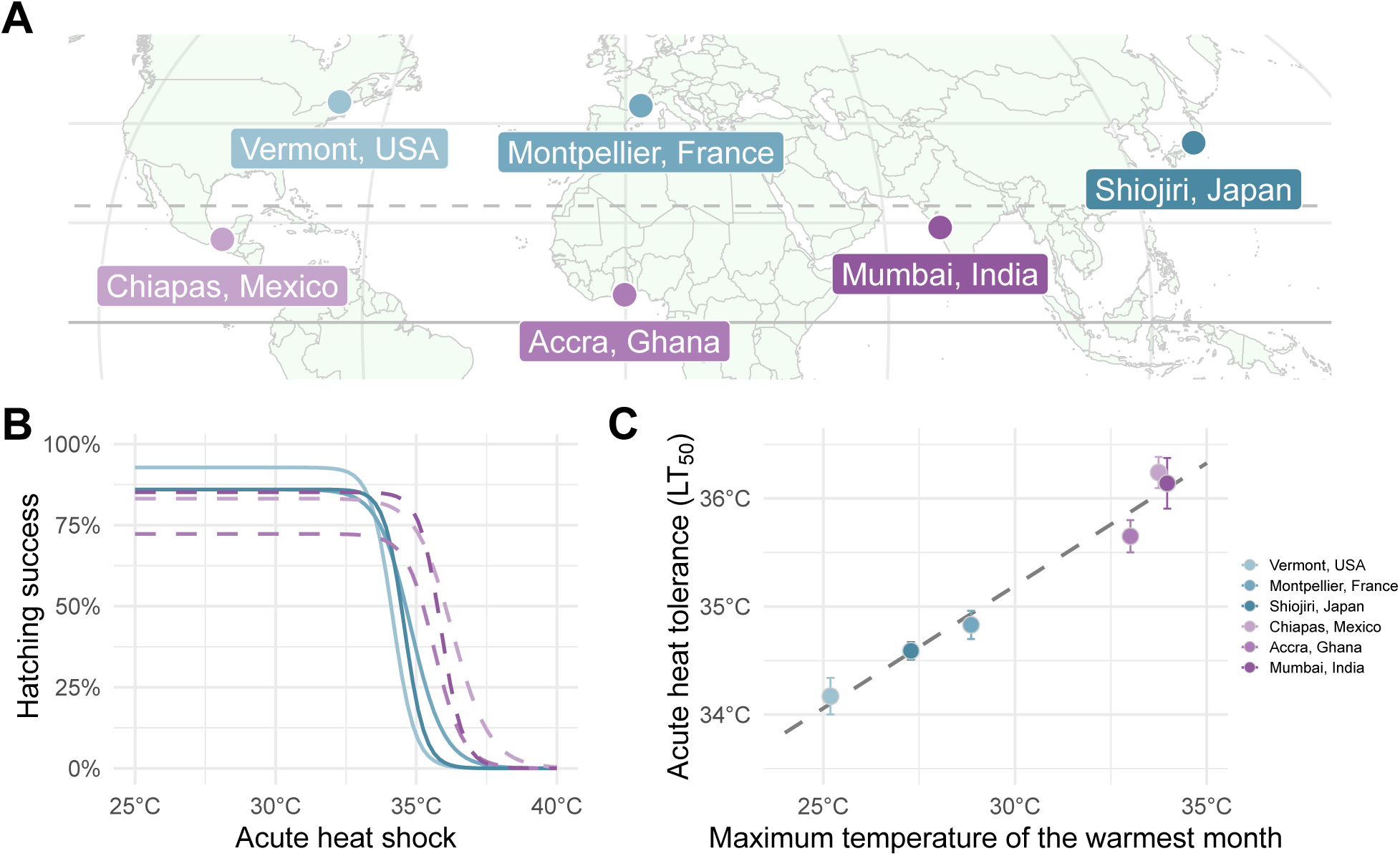
Acute heat tolerance of early embryos is higher in warmer climates. (**A**) Map with the collection locations for each genotype. (**B**) Hatching success nonlinear survival curve versus acute heat shock temperature. (**C**) Scatter plot of embryonic acute heat tolerance (LT_50_) versus the maximum temperature of the warmest month (Bioclimatic variable 5; WorldClim) for each collection location.

### Metabolomics and data normalization

Using untargeted liquid chromatography–mass spectrometry (LC–MS), we initially detected 45 metabolites of interest. After applying quality control filters to remove metabolites with unreliable quantification, 33 metabolites remained for downstream analysis (**Fig. S1A**). We also calculated the ratios of three major redox couples — NADH/NAD^+^, NADPH/NADP^+^, and reduced and oxidized glutathione (GSH/GSSG) — and included these ratios in all subsequent analyses. Pareto scaling of metabolite intensities yielded comparable intensity distributions across all samples, indicating effective normalization (**Fig. S1B**).

### Principal component and clustering analyses

Principal component analysis (PCA) including the pooled quality control (QC) samples confirmed minimal technical variation in the LC–MS protocol, with all QC samples clustered tightly near the origin (**Fig. S2A**). Overall, PCA revealed a marked shift in the metabolites following heat shock, with the samples generally moving down PC2 and to the right on PC1 (**Fig. 2A**). 54.0% of the variance was explained by the first PC, with the second PC explaining 15.9% (92.2% explained by the first 6 PCs; **Fig. S2B**). All metabolites showed positive loadings on PC1, indicating a concerted shift in overall metabolite abundance following heat shock (**Fig. S2C**). Hierarchical clustering (Pearson correlation) separated the metabolite profiles into three major groups (**Fig. 2B**). The first cluster (n = 26) comprised exclusively heat-shocked embryos (all 18 temperate and 6 tropical), defining a group of heat-responsive samples. The second cluster (n = 23) was dominated by tropical samples at 25°C (18; 82%), alongside a minority of heat-shocked tropical samples and a single temperate control sample, reflecting mostly the baseline tropical metabolome, and heat-insensitive tropical samples. The third cluster (n = 23) consisted mainly of temperate embryos at 25°C (17; 74%), with a subset of heat-shocked tropical samples (6; 26%) possibly indicative of tropical samples with a moderate response to heat shock. Together these results highlight a robust metabolic response to heat shock in the temperate metabolome, alongside a more heterogeneous response in the tropical embryos. Notably, the heterogeneous response in tropical embryos did not segregate by genotype. Each tropical genotype contributed heat-shocked replicates to multiple clusters: Chiapas, Mexico replicates were split evenly between clusters 1 and 2, Mumbai, India replicates were split evenly between clusters 2 and 3 and Accra, Ghana had five replicates in cluster 1 and one replicate in cluster 2. Therefore, no tropical genotype formed a distinct cluster of heat-shocked samples, underscoring the within-genotype variability in the metabolomic response to heat.

**Figure 2.**
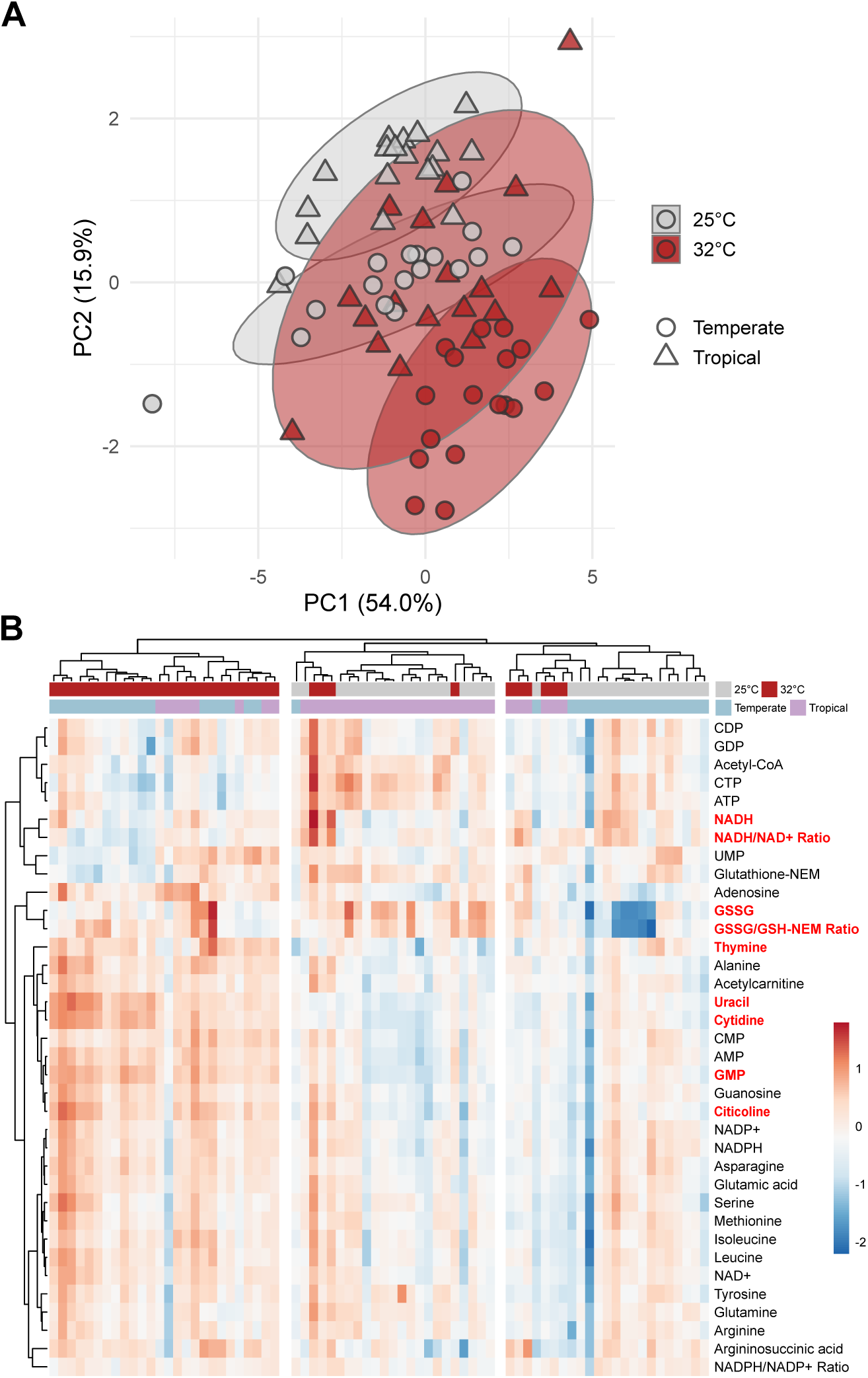
Heat shock drives a region-specific metabolomic shift with greater post-shock variance in tropical embryos. (**A**) Scatter plot of PC1 versus PC2. Points are colored by treatment: grey points are 25°C, red points are heat-shocked; shape indicates region of origin: temperate samples are circles, and tropical samples are triangles. Ellipses indicate a 95% confidence region for the treatment-region group. (**B**) Heatmap of the 33 metabolites and 3 redox ratios (rows) across the 72 samples (columns). The rows and columns are clustered by their Pearson correlation. Metabolites and ratio names highlighted in red indicate a significant temperature and region interaction (adj. p < 0.05). Color scale reflects relative abundance (blue is low, white is medium, and red is high).

### Metabolomics PCs correlate with embryonic heat tolerance

Next, we examined whether overall metabolome profiles, as summarized by principal components, predict LT_50_ under control (25°C) and heat shock (32°C) conditions. Of all PCs, only PC2 and PC4 showed significant associations with LT_50_ across both treatments. PC2 scores increased with increasing LT_50_ in both 25°C and 32°C treatments (**Fig. 3A**; 25°C: slope = 0.67, S.E. = 0.12, R^2^ = 0.49, adj. p = 0.00017; 32°C: slope = 0.81, S.E. = 0.18, R^2^ = 0.37, adj. p = 0.0018). PC2 was dominated by positive loadings of CTP and ATP (contributing 17.8% and 9.1% to the PC2 axis, respectively), indicating that this PC reflects nucleotide energy reserves (**Fig. 3B**). In both control and heat-shocked samples a higher PC2 score was correlated with greater heat tolerance, reflecting increases in ATP and CTP. In contrast, PC4 displayed a treatment dependent relationship with heat tolerance (**Fig. 3C**; 25°C: slope = 0.35, S.E. = 0.08, R^2^ = 0.39, adj. p = 0.0015; 32°C: slope = –0.45, S.E. = 0.13, R^2^ = 0.26, adj. p = 0.020) and was driven primarily by the ratio of NADH to NAD^+^ (**Fig. 3D**; contributing 21.8% to PC4). At 25°C increases in PC4 were indicative of greater heat tolerance, whereas at 32°C increased heat tolerance was negatively correlated with PC4 scores.

**Figure 3.**
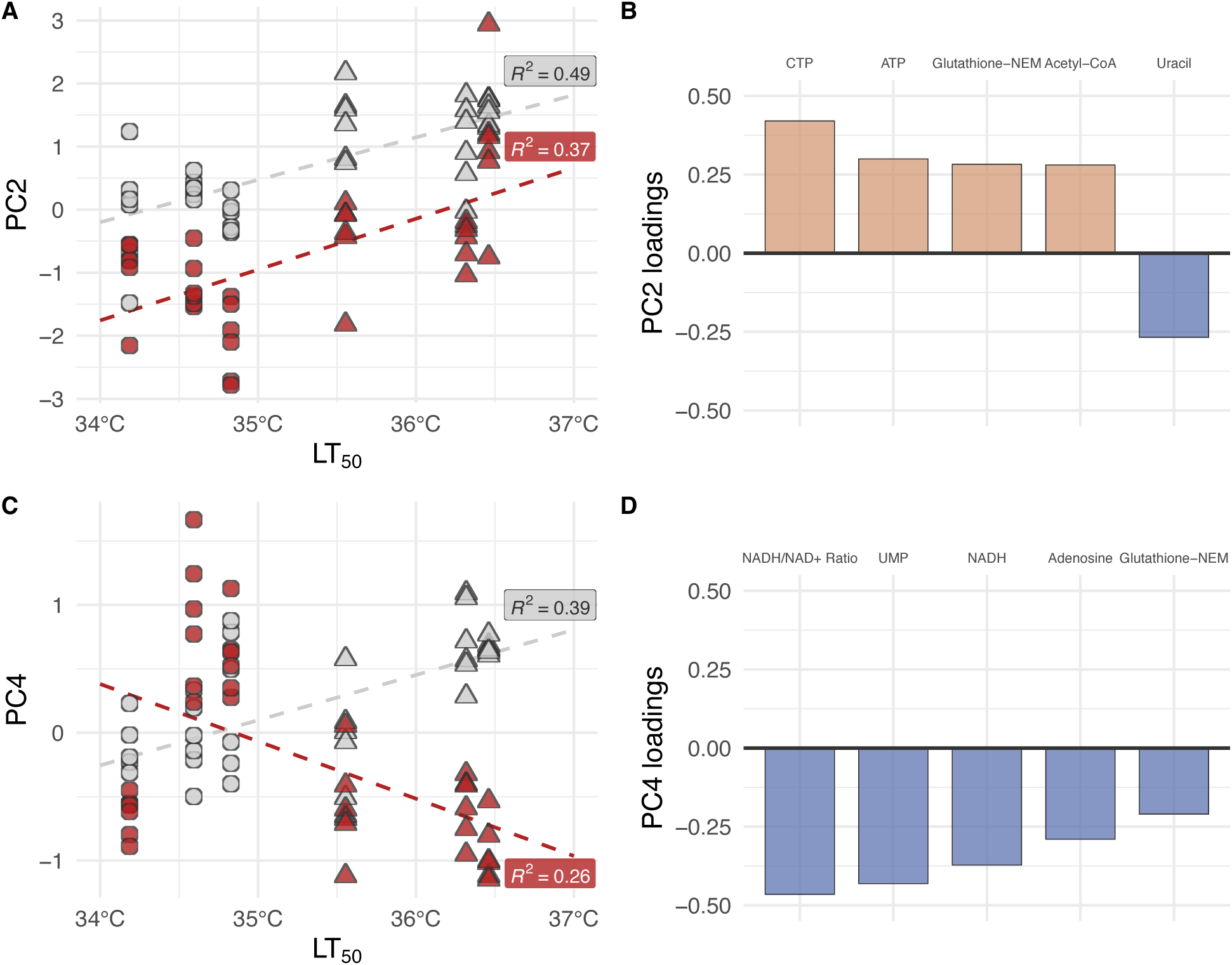
Embryonic heat tolerance (LT_50_) increases with PC2 at 25°C and after heat shock, whereas PC4 shows a treatment-specific association with LT_50_. (**A**) Scatter plot of LT_50_ versus PC2 scores for each genotype at baseline (25°C, grey) and after heat shock (32°C, red). Points are shaped by region (temperate: circle; tropical: triangle) and dashed lines show separate linear model fits for each treatment. (**B**) Bar plot showing the five metabolites with the largest absolute loadings on PC2 (from the PCA rotation matrix). Bars are colored by loading direction: positive loadings in orange and negative loadings in blue. (**C**) Scatter plot of LT_50_ versus PC4 scores as in (A). (**D**) Bar plot of the top five metabolites with the largest absolute loadings on PC4 derived from the PCA rotation matrix.

### Differentially abundant metabolites

Using linear mixed-effects models to evaluate temperature and region effects, we detected no significant main effect of region on any metabolite (**Fig. S3A**; adj. p < 0.05). In contrast, a majority of the metabolites (66.7%; n = 24) responded to acute heat shock, with 23 of 24 (96%) showing an increase following heat shock (**Fig. S3B**; adj. p < 0.05), underscoring the predominately shared metabolomic response between regions. However, nine metabolites and ratios (GMP, uracil, thymine, cytidine, citicoline, GSSG, NADH, NADH/NAD^+^ and GSH/GSSG) displayed a significant interaction effect between temperature and region (adj. p < 0.05). The region-specific differences were largely in the magnitude of change and not direction, with only four redox-related metabolites and ratios showing a difference in directionality of change between regions (**Fig. S3C**). Specifically, GMP, uracil, thymine, cytidine, and citicoline all showed a greater heat-induced increase in temperate embryos than tropical embryos (**Fig. S4**), showing that among these non-redox related metabolites, region-specific response was in the magnitude of response. However, the redox-related metabolites and ratios (NADH, GSSG, NADH/NAD^+^, and GSH/GSSG) showed a distinct region-specific pattern with the temperate embryos becoming more oxidized, and tropical embryos either maintaining redox status or becoming slightly more reduced (**Fig. S4**). After heat shock, NADH increased in abundance by 2.2-fold in tropical embryos (post hoc temperature contrast within region, adj. p = 0.0059) but did not change in temperate embryos (**Fig. S5**; adj. p = 0.55). Meanwhile, oxidized glutathione (GSSG) increased by 4-fold in temperate embryos after heat shock (adj. p = 0.049) but did not significantly change in tropical embryos (**Fig. S5**; adj. p = 0.15).

### Region-specific shift in redox couple ratios

Both the NADH/NAD^+^ and GSH/GSSG showed a region-specific response to temperature (**Fig. 4, A and C**; region × temperature, adj. p = 0.0005 and 0.011, respectively). After heat shock, in temperate embryos NADH/NAD^+^ ratio declined by 46.6% to a more oxidized state (post hoc temperature contrast within region, adj. p = 0.0035), while in tropical embryos the ratio increased by 52.9% to a more reduced state (adj. p = 0.0058). NADPH/NADP^+^ ratio did not display a significant region-specific response to temperature (adj. p = 0.16); although, the median values trended in the same direction, with temperate lines showing greater oxidation and tropical lines greater reduction or little redox change following heat stress (**Fig. 4B**). Parallel changes in the absolute abundances of reduced versus oxidized forms of these metabolites support this region-specific redox shift (**Fig. S5**). At the individual genotype level, heat-tolerant lines generally became more reduced (or had little change in redox status), following heat shock while heat-sensitive lines became more oxidized (**Fig. S6, A–C**), highlighting the link between redox status and embryonic heat tolerance.

**Figure 4.**
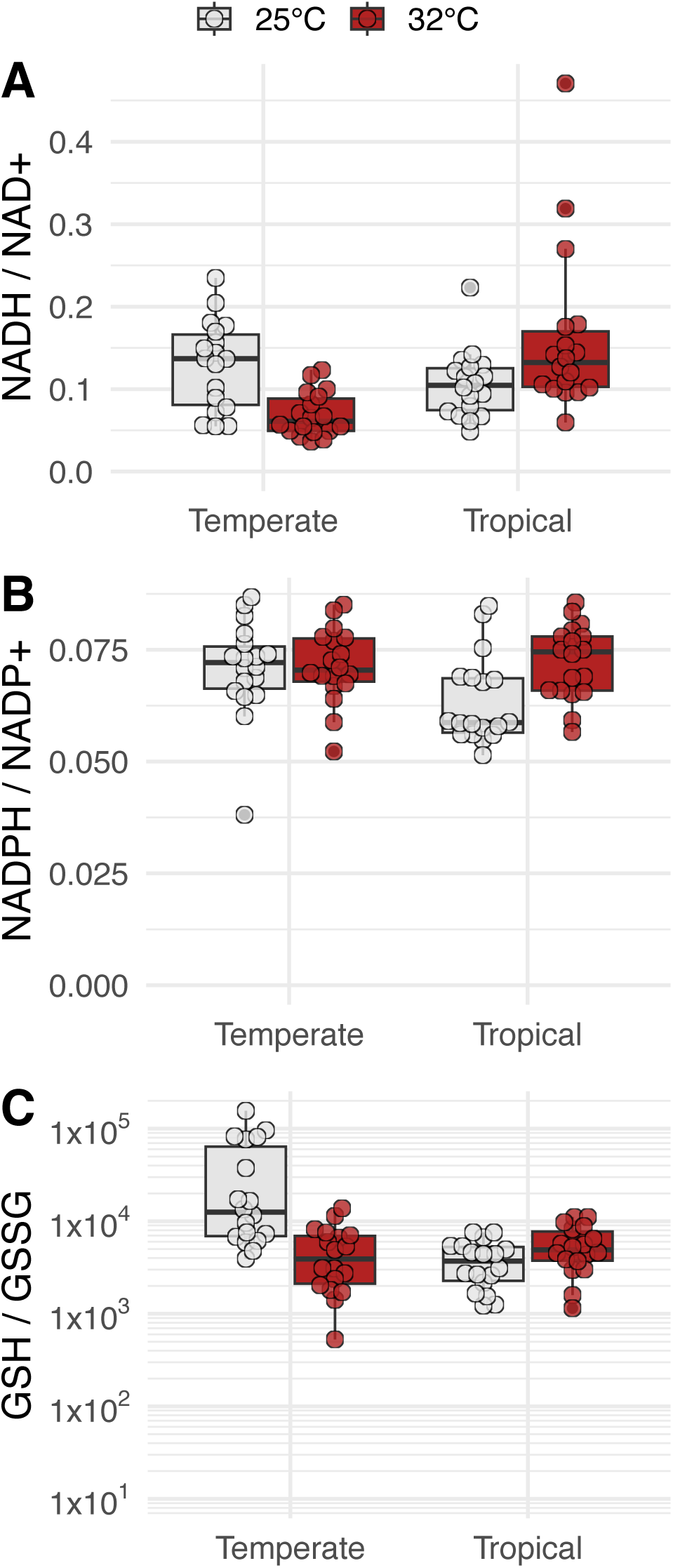
Major redox couples shift in opposite directions following acute heat shock: temperate embryos become more oxidized and tropical embryos become more reduced. Box plots of the ratios of reduced to oxidized form of the redox couples: NADH/NAD^+^ (**A**), NADPH/NADP^+^ (**B**), and reduced glutathione (GSH) / oxidized glutathione (GSSG) (**C**). Boxes represent the interquartile range, middle bar indicates the median, whiskers extend to 1.5×IQR. Overlaid individual points represent pools of 1,000 embryos (N = 72).

### Changes in nucleotide monophosphates following acute heat shock

Three nucleotide monophosphates — AMP, CMP, and GMP — increased in abundance following 32°C heat shock in both temperate and tropical embryos (**Fig. 5, A–C**; adj. p = 1.1 × 10^−11^, 0.00014, and 1.6 × 10^−23^ respectively). In AMP and CMP, both regions responded similarly (**Fig. 5, A and B**; interaction region × temperature adj. p > 0.05). However, in temperate embryos, GMP showed a greater response to heat stress with a 2.89-fold increase in abundance, compared to 2.32-fold increase in tropical embryos (**Fig. 5C**; interaction region × temperature adj. p = 0.007).

**Figure 5.**
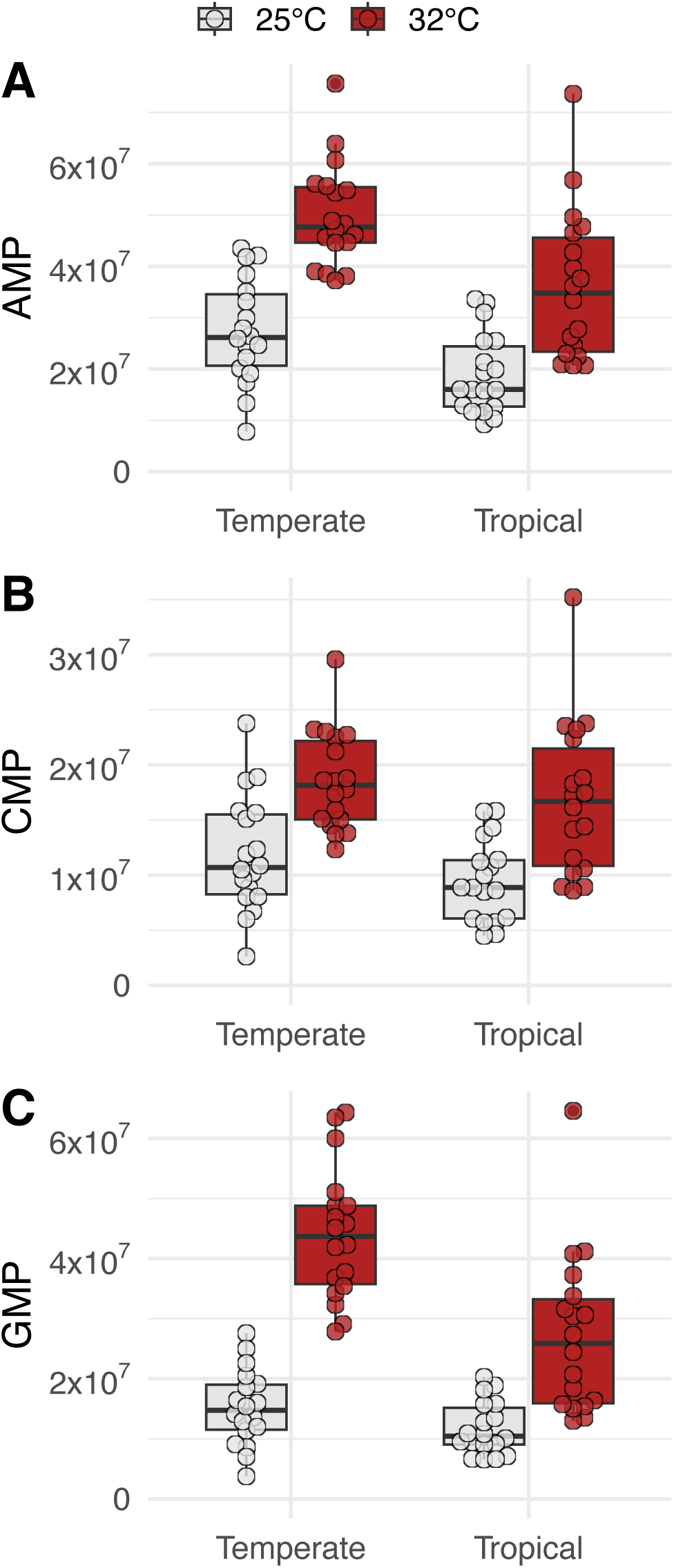
Heat shock increases the abundance of nucleotide monophosphates in both temperate and tropical embryos. Box plots of the normalized intensity of three nucleotide monophosphates: AMP (**A**), CMP (**B**), and GMP (**C**). Boxes represent the interquartile range, middle bar indicates the median, whiskers extend to 1.5×IQR. Overlaid individual points represent pools of 1,000 embryos (N = 72).

## Supplementary materials

**Table S1.** Fly stock numbers, collection location, and embryonic LT_50_.

| Location | Stock No. | Latitude | Longitude | Embryonic LT<sub>50</sub> |
| --- | --- | --- | --- | --- |
| Vermont, USA | VTECK10 | 44.37 | -72.43 | 34.2°C |
| Montpellier, France | 14021-0231.149 | 43.61 | 3.88 | 34.8°C |
| Shiojiri, Japan | 14021-0231.135 | 36.11 | 137.95 | 34.6°C |
| Chiapas, Mexico | 14021-0231.22 | 16.70 | -93.01 | 36.3°C |
| Accra, Ghana | 14021-0231.182 | 5.56 | -0.20 | 35.6°C |
| Mumbai, India | 14021-0231.45 | 19.08 | 72.88 | 36.5°C |

**Figure S1.**
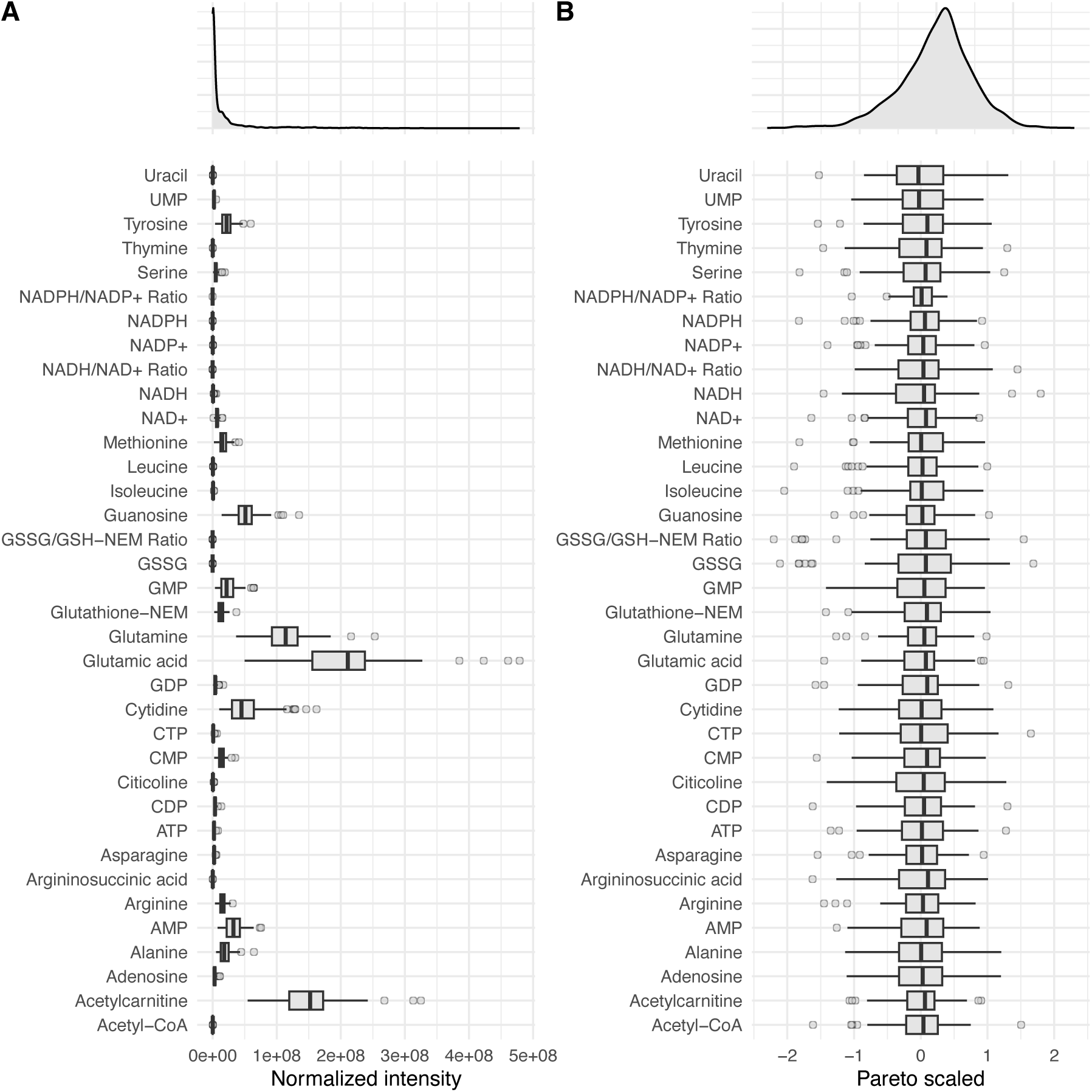
Normalization of metabolomics data. (**A**) Top: density plot of normalized intensity across all metabolites. Bottom: box plot of normalized intensity for all quantified metabolites. (**B**) Top: density plot of pareto scaled data across all metabolites. Bottom: box plot of same data pareto scaled. Boxes represent the interquartile range, middle bar indicates the median, whiskers extend to 1.5×IQR and individual samples beyond 1.5×IQR are plotted individually.

**Figure S2.**
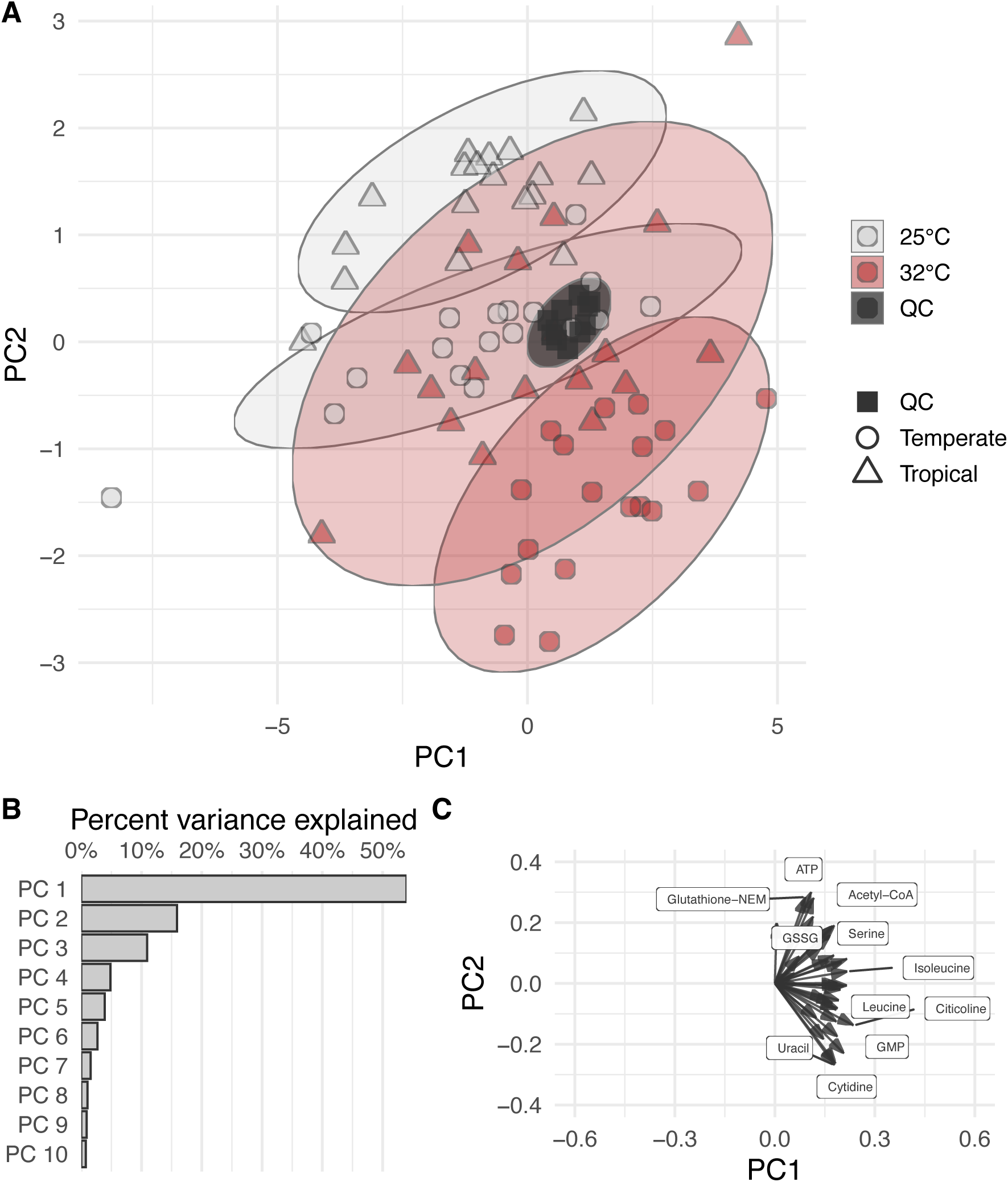
Principal component analysis of the metabolite abundances. (**A**) Scatter plot of PC1 versus PC2. Points are colored by treatment: grey points are 25°C, red points are 32°C; shape indicates region of origin: temperate samples are circles, and tropical samples are triangles. Quality control samples are dark grey boxes. Ellipses indicate a 95% confidence region for the treatment-region group. (**B**) Scree plot of the percent variance explained by the first 10 principal components. (**C**) Rotation (loading) plot showing each metabolite’s contribution to PC1 and PC2. Arrows originate at the origin and point to the PC1 and PC2 loading coordinates for each compound; the length of each arrow reflects the magnitude of that metabolite’s contribution.

**Figure S3.**
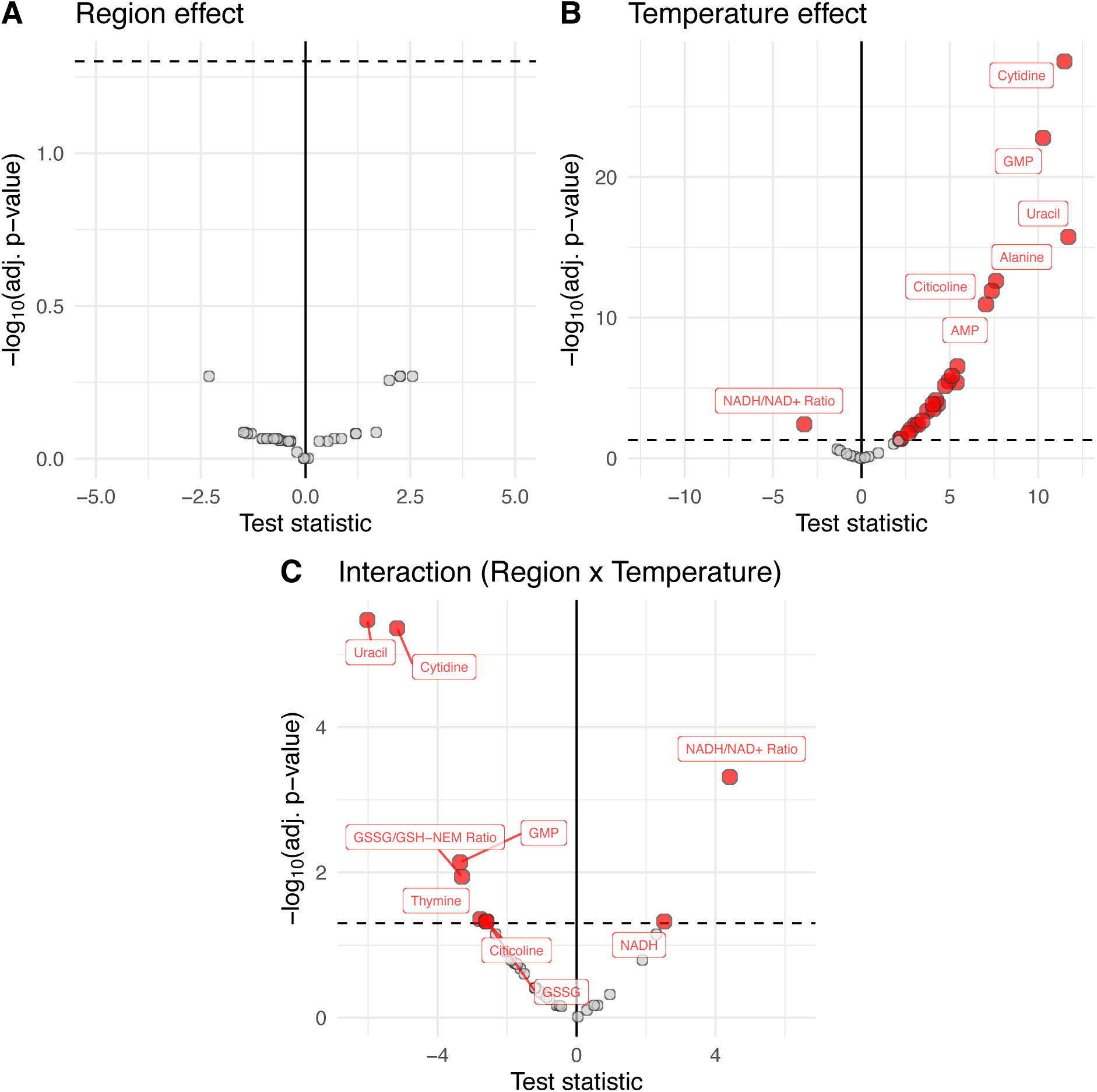
Results of the differentially abundant metabolites show significant temperature and temperature × region interaction effects. Volcano plots of the linear-mixed effects model results with the test statistic (horizontal axis) versus -log_10_(adj. p-value) (vertical axis) for the region effect (**A**), temperature effect (**B**) and, (**C**) interaction between region and temperature. Points are colored by significance: not-significant (grey), significant (red). Dashed horizontal line indicates the significance threshold (adj. p-value < 0.05).

**Figure S4.**
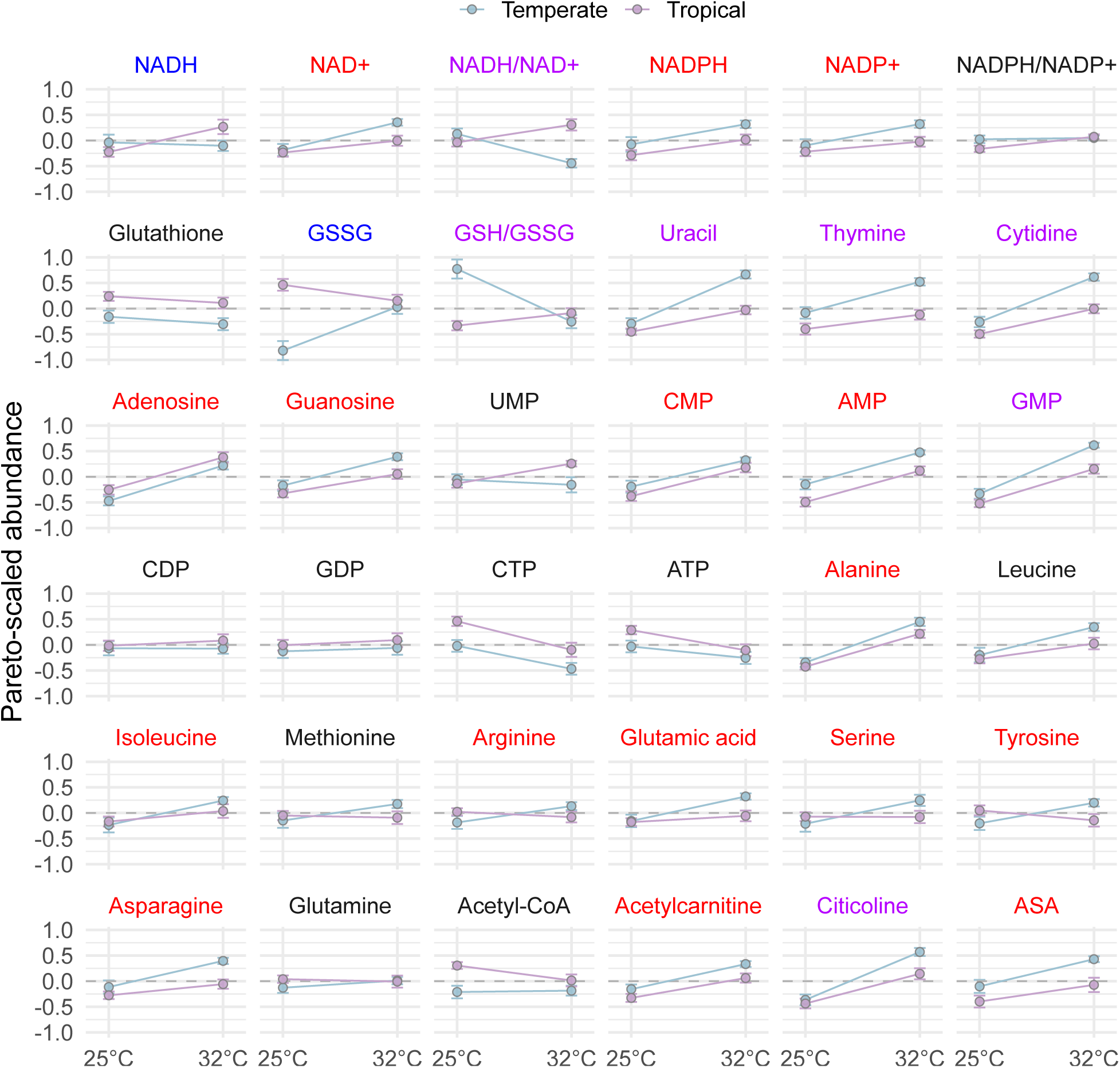
Reaction norm plots for all 33 quantified metabolites and 3 redox ratios. Points represent the mean of the pareto-scaled abundance for temperate and tropical embryos with error bars indicating the standard error. Metabolite names are colored by significance of the differential abundance testing; red indicates a significant effect of temperature, purple indicates both a significant effect of temperature and a significant interaction between temperature and region, and blue indicates only a significant interaction between temperature and region.

**Figure S5.**
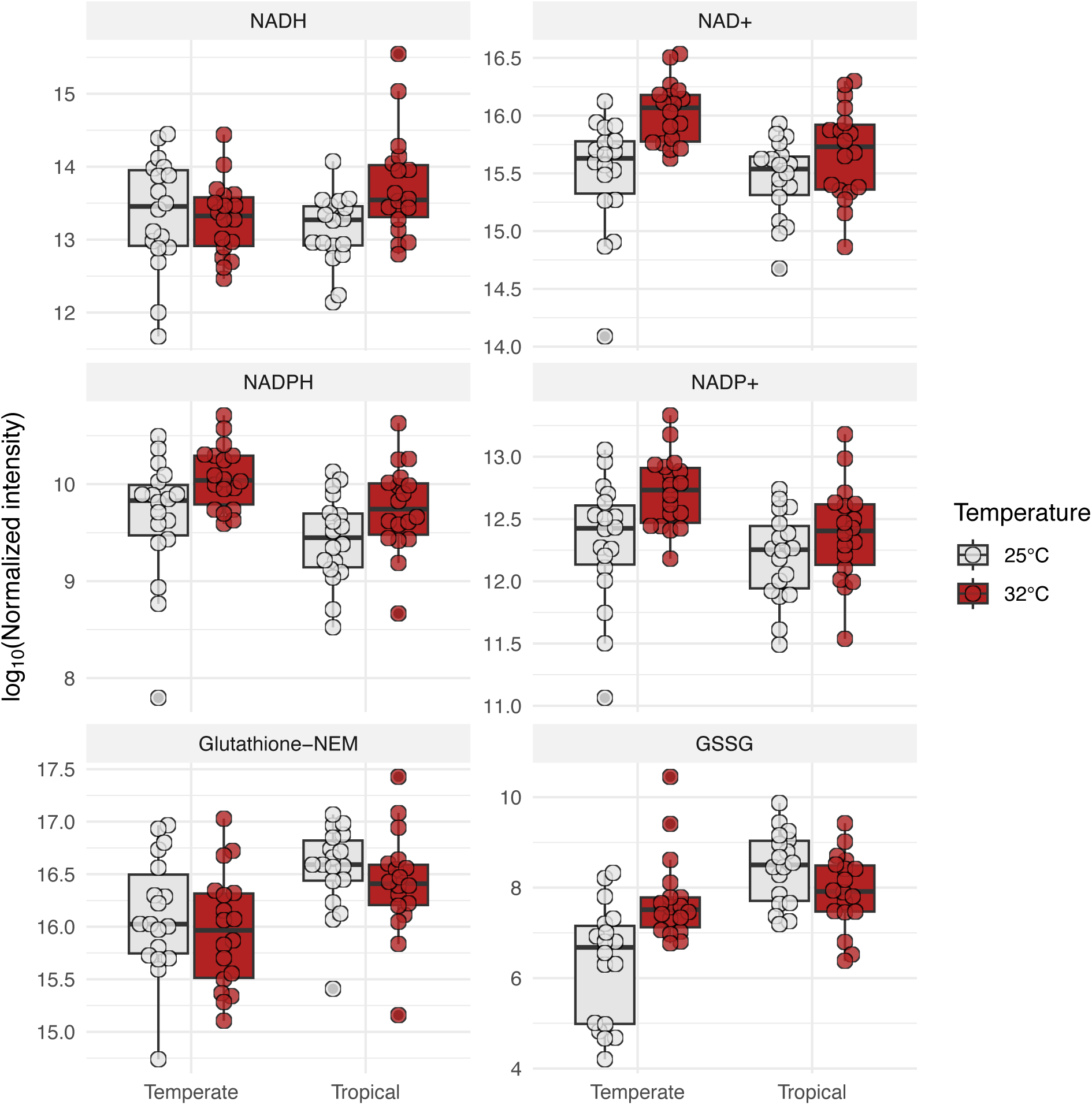
Response to heat shock of individual reduced and oxidized forms of the redox couples. Box plots of the log_10_ normalized intensity of the reduced and oxidized form of three major redox couples. Boxes represent the interquartile range, middle bar indicates the median, whiskers extend to 1.5×IQR. Overlaid individual points represent pools of 1,000 embryos (N = 72).

**Figure S6.**
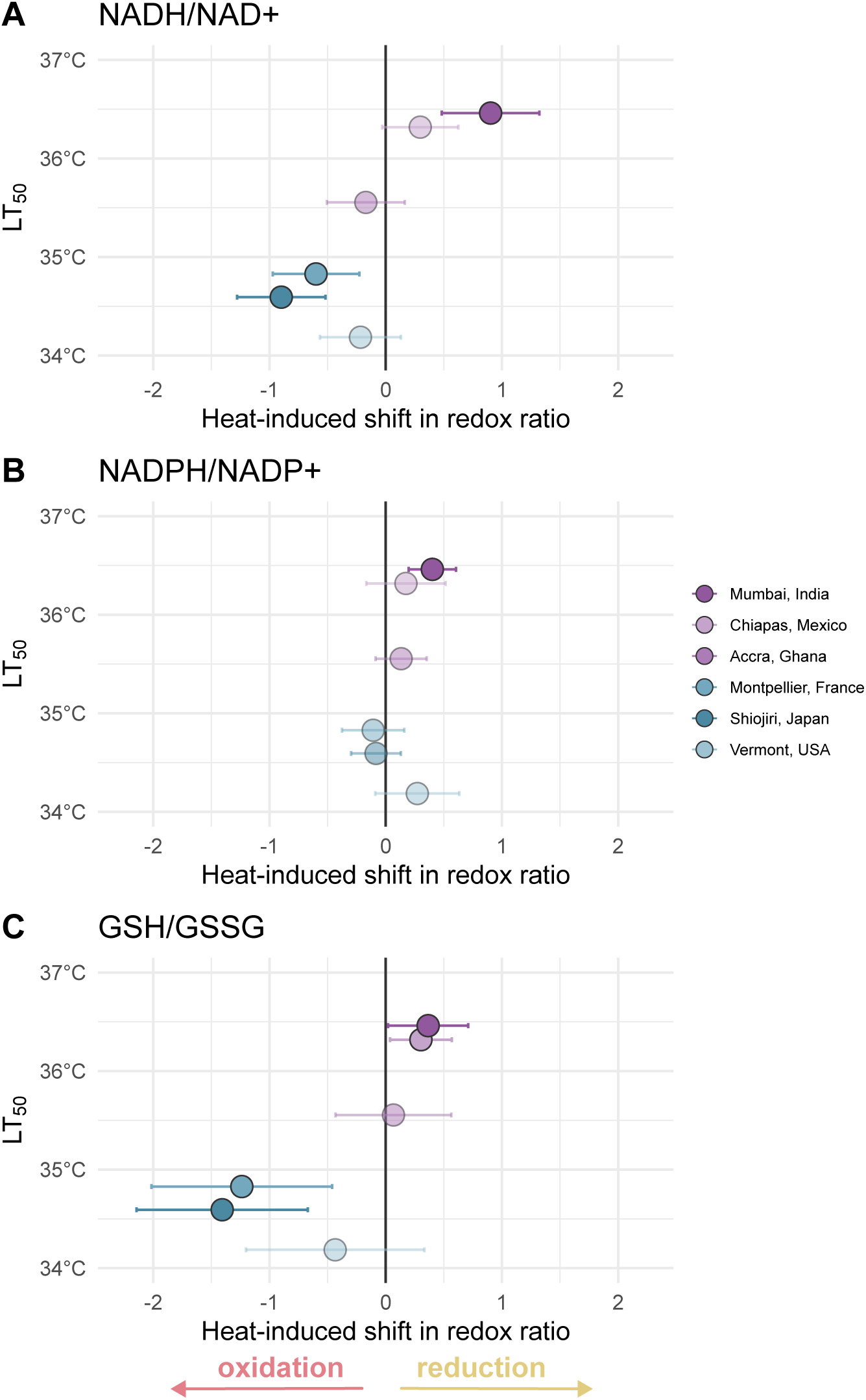
Redox state shift in major redox couples correlates loosely with embryonic heat tolerance. Mean changes in pareto-scaled redox ratios (32°C – 25°C ± 95% CI) versus LT_50_ for (**A**) NADH/NAD^+^, (**B**) NADPH/NADP^+^, and (**C**) GSH/GSSG. Points are colored by genotype. Points to the right of zero indicate a heat induced shift towards reduction, whereas shifts to the left indicate a move towards a more oxidative state; points with 95 % CIs overlapping zero are included at reduced opacity.

## Discussion

Acute heat shock induced rapid and largely shared metabolic changes in early *D. melanogaster* embryos of both temperate and tropical regions, but the response diverged among redox metabolites, where temperate embryos shifted towards an oxidized state, and tropical embryos either maintained redox balance or shifted to a reduced state. Although temperate and tropical embryos both showed increased energy demand and nucleotide turnover after heat stress, the downstream effects seem to be distinct. Temperate embryos showed more evidence of redox imbalance and possibly ROS production after heat shock, while tropical embryos displayed a more varied response, with some replicates and metabolites considerably less affected by heat stress. These region-specific redox shifts align with the acute heat tolerance phenotype, with temperate embryos more heat sensitive and tropical embryos more heat tolerant (Lockwood et al., 2018). Taken together, these patterns support the hypothesis that maintaining tight control of redox balance is essential during embryonic development, especially when dealing with heat stress (Mikucki et al., 2024; Petrova et al., 2018; Takahashi, 2012; Timme-Laragy et al., 2013; Ufer and Wang, 2011).

Across regions, this largely shared metabolomic response to heat shock — with increases in nucleotide monophosphates (*e.g.*, AMP, CMP, and GMP), pyrimidines bases/nucleosides (*e.g.*, uracil, thymine, cytidine), and amino acids (*e.g.*, alanine, glutamic acid, serine) — is consistent with increased biochemical kinetics due to *Q*_10_ effects (Arrhenius, 1889; Kohout, 2021; Somero et al., 2017) and a heat-induced proteostasis response, with ATP-intensive chaperone activity and increased free amino acids following proteolysis of damaged proteins (Feder and Hofmann, 1999; Laufen et al., 1999; Malmendal et al., 2006; Somero, 1995). These patterns align with previous studies which have shown that acute heat stress (*i.e.*, minutes to hours) can rapidly shift metabolite pools linked to ATP production (Malmendal et al., 2006; Xu et al., 2025), and other studies which show that even moderate differences in the thermal environment can alter carbohydrate and lipid energy stores (Klepsatel et al., 2016; MacMillan et al., 2016; Sarup et al., 2016; Schou et al., 2017). These rapid metabolic changes mirror the robust transcriptomic response observed across similar genotypes of *D. melanogaster* embryos, where the response was largely shared across regions with the notable exception of the genes related to oxidative stress (Mikucki et al., 2024). Echoing this pattern, here the redox-related metabolites represent an exception to the shared heat stress response.

After heat shock, the opposing directional shifts in NADH/NAD^+^ ratio are consistent with region-specific differences in the relative rates of NADH production and oxidation (Vinogradov, 2008; Xiao et al., 2018). Temperate embryos showed a post–heat-shock decrease in NADH/NAD^+^ ratio, while tropical embryos showed an increase accompanied by higher NADH abundance. Although we did not measure flux, this pattern is consistent with temperate embryos oxidizing NADH more rapidly *via* the electron transport system (ETS), while tropical embryos generate NADH faster than it is oxidized under heat stress (Adam-Vizi and Chinopoulos, 2006; Hung et al., 2011). This imbalance is significant because sustained ATP generation requires continual re-oxidation of NADH and the shuttling of reducing equivalents (*e.g.*, malate-aspartate shuttle) to the ETS to power oxidative phosphorylation (Somero et al., 2017), so a mismatch between production and oxidation is expected to slow ATP output, bottleneck electrons on the ETS, and increase ROS formation (Zhao et al., 2019). This can have particularly detrimental consequences on embryogenesis, as early embryos rely on oxidative phosphorylation (Tennessen et al., 2014) and even transient loss of mitochondrial ATP production is known to disrupt nuclear divisions in the syncytial embryo (Chowdhary et al., 2017).

Region-specific differences are also captured in the principal component analysis at both control and heat-shock temperatures. Although we did not detect a significant effect of region for any individual metabolite, regional differences in redox metabolites and energy stores are apparent with PC2 scores (driven largely by ATP, CTP, and reduced glutathione) correlated with increased embryonic heat tolerance. PCA is well-suited to capture these coordinated, low effect-size differences across metabolites that the linear-mixed effects model is less able to detect (Worley and Powers, 2013). Overall, this pattern suggests that heat-tolerant embryos maintain higher energy reserves and reducing equivalents at 25°C, which may provide these embryos with a buffer against heat-induced energy depletion and oxidative shift. Beyond these effects of ATP supply and redox balance, shifts in cellular redox state can also affect posttranslational modifications and disrupt developmental progression. During the oocyte-to-embryo transition, proteome-wide shifts in thiol reactivity show the redox sensitivity of key developmental regulators; perturbing cellular redox state at this stage disrupts cell-cycle timing and likely influences chromatin regulation and translation (Petrova et al., 2018). A shift in thiol reactivity would be expected to be mirrored in the glutathione ratio (GSH/GSSG) (Go and Jones, 2013; Schafer and Buettner, 2001).

Heat shock produced region-specific shifts in glutathione redox balance — a key marker of oxidative stress (Forman et al., 2009; Lu and Holmgren, 2014; Schafer and Buettner, 2001) — with temperate embryos shifting toward higher oxidation (increased GSSG relative to GSH), while tropical embryos maintained a stable GSH/GSSG ratio. The glutathione pathway functions as a two-step cycle where (i) glutathione-dependent peroxidases reduce peroxides (*e.g.*, H_2_O_2_) to water while oxidizing two molecules of GSH, and subsequently (ii) glutathione reductase (GR) uses NADPH to reduce oxidized glutathione back to GSH (Maiorino et al., 2007; Radyuk et al., 2010; Singh et al., 2001; Tu and Akgül, 2005; Vašková et al., 2023). The oxidized shift in temperate embryos is consistent with increased ROS production during thermal stress, a known consequence of high metabolic rates at warm temperatures (Banh et al., 2016; Chung and Schulte, 2020; Farahani et al., 2020; Hasanuzzaman et al., 2020; Rossi et al., 2024; Zhu et al., 2017). However, because glutathione balance is maintained in a cycle, the stable GSH/GSSG ratio in tropical embryos does not preclude ROS production, and it may instead suggest more effective recycling capacity (Forman et al., 2009). The lack of change in the NADPH/NADP^+^ ratio in both regions is likely reflective of the many roles of NADP(H) (*e.g.*, pentose-phosphate pathway) and may not solely relate to its actions within the glutathione pathway (Stincone et al., 2015; Ying, 2008).

The known mechanisms of mitochondrial ROS formation suggest the possibility of heat-induced ROS accumulation in both temperate and tropical embryos. In a reduced state with a high NADH/NAD^+^ ratio, as observed in the heat-shocked tropical embryos, ROS production is most likely to occur *via* reverse electron transport (RET), where complex I transports electrons back to form superoxide; but this occurs only in combination with a high proton gradient and a reduced coenzyme Q pool (Murphy, 2009). By contrast, when NADH is being rapidly oxidized, as in temperate embryos under heat stress, ROS is expected to form at several sites primarily *via* forward electron leak in complex I (flavin), mitochondrial glycerol-3-phosphate dehydrogenase (mGPDH), and complex III (Q_o_) (Brand, 2010; Graham et al., 2022; Miwa and Brand, 2005; Miwa et al., 2003; Murphy, 2009; Treberg et al., 2011). Overall, these patterns (*i.e.*, faster NADH oxidation and increased oxidized glutathione) suggest higher ETS flux and ROS formation in temperate embryos, in contrast with better glutathione recycling in tropical embryos. These predictions could be tested with a variety of assays, including enzymatic rate measurements (*e.g.*, GR activity), mitochondrial respiration assays (*e.g.*, Seahorse), compartment-specific ROS probes (*e.g.*, MitoSOX) and NADH/NAD^+^ sensors (Hung et al., 2011; Jørgensen et al., 2021; Mannervik, 2001; Mukhopadhyay et al., 2007).

Finally, the relatively heterogeneous tropical response to heat shock likely reflects a nonlinear thermal response, with our heat-shock temperature near a critical threshold for redox balance (Huey and Kingsolver, 1989; Miwa and Brand, 2005; Murphy, 2009; Somero et al., 2017; van Heerwaarden et al., 2024). Under this model, temperate embryos have largely exceeded this tipping point and consequently lost redox balance, while tropical embryos are straddling it, with some maintaining balance and some beginning to transition to a stress-induced imbalance. This pattern is evident across redox and non-redox metabolites, with several tropical heat-shocked replicates clustering with the 25°C controls, while others closely align to the heat-shocked temperate metabolomic profiles. Overall, this result mirrors the acute heat shock phenotype where survival rates plummet after a critical threshold (Lockwood et al., 2018). Together, these findings highlight that maintaining redox balance may be an underappreciated determinant of acute heat tolerance, especially in rapidly developing early embryos.

## Methods

### Fly care

For several generations prior to experimentation, we maintained fly lines on standard yeast, cornmeal, and molasses food at a standard density of approximately 50 to 100 flies per vial (95 mm × 25 mm, Genesee Scientific). We maintained the lines at 25°C and 55% relative humidity on a 12hr:12hr light:dark cycle (DR-36VL, Percival Scientific Inc.). We sourced isofemale fly lines from the National *Drosophila* Species Stock Center at Cornell University (Montpellier, France; Shiojiri, Japan), the UC San Diego *Drosophila* Species Stock Center (Chiapas, Mexico; Accra, Ghana; Mumbai, India), and from wild-caught females in East Calais, Vermont (Cooper et al., 2014) (**Table S1**).

### Acute heat tolerance assay

Prior to embryo collection, we prepared small fly cages (Genesee Scientific) with 1–2 day-old adult flies with grape juice agar plates (60 mm × 15 mm) with yeast paste. We incubated the cages at 25°C and replaced the agar plates and yeast paste every 24 hours. Two hours before embryo collection, we supplied each cage with a fresh agar plate with yeast to prevent egg retention. To collect embryos, we exchanged the pre-lay plate with a new grape-juice agar plate with yeast paste and allowed the flies to lay eggs for one hour. Immediately following collection, the egg plates were wrapped in parafilm and submerged in a water bath for 45 minutes at temperatures ranging from 20°C to 44°C. After heat shock, we counted the embryos and placed them into neat grids with a paintbrush for hatching assessment. We recorded the proportion of eggs hatched 48 hours after egg laying.

### Egg collection & heat shock for metabolomics

For each genotype, we set up 8–10 cages, each containing 100 mating pairs to accommodate the large number of embryos needed for quality quantification in the metabolomics experiment. We collected embryos and treated them to 25 or 32°C as described above. Immediately after heat shock, we washed the embryos in 0.7% (*w/v*) NaCl, 0.05% Triton X-100, rinsed with dH_2_O, and transferred the clean embryos to Whatman paper to dry while counting out 1,000 embryos. We then transferred the embryos into PowerBead tubes (Qiagen) containing 1.4 mm ceramic beads, froze the samples in liquid nitrogen, and stored them at –80°C. For each group, we collected 6 replicates, yielding a total of 72 samples across 6 genotypes and two temperatures.

### Sample preparation for metabolomics

All metabolomics and mass spectrometry was completed in the Metabolomics Core Facility at the University of Utah in Salt Lake City, Utah. We shipped the samples overnight on dry ice. Following the core facility’s standard protocol, they extracted each 1,000-embryo sample in a solution of ice-cold 14:1:5 ACN:MeOH:H_2_O containing 0.1 % NH_4_OH, 0.1 µg/mL D_9_-carnitine, 1.25 µM Metabolomics Amino Acid Mix Standard (Cambridge Isotope Laboratories, Inc, Andover, MA, USA), and 1.5 mg/mL N-ethylmaleimide as a derivatization reagent for free glutathione. A process blank containing only the extraction solution was created and carried throughout the extraction. Each sample was homogenized in bead mill tubes with ceramic beads for 30 seconds prior to incubation at –20°C for 1 h. The samples were then transferred to new 1.7 mL Eppendorf tubes and centrifuged at 20,000 × g for 10 minutes at 4°C. The supernatant was transferred to new PTFE autosampler vials for immediate analysis.

### Mass spectrometry of samples

Pooled quality control (QC) samples were prepared by combining 10 µL of each sample, and the sample injection order was randomized prior to analysis. Mass spectrometry was performed at the Metabolomics Core Facility using a SCIEX 7600 Zeno-ToF (AB SCIEX LLC, Framingham, MA, USA) coupled to an Agilent 1290 Infinity II HPLC system in positive-ionization mode. Chromatographic separation was achieved on a Waters BEH Z-HILIC 2.1 × 100 mm column (Waters Corporation, Milford, MA, USA) with Phenomenex Krudkatcher Ultra (Phenomenex, Torrance, CA, USA). Chromatography was performed using buffer A (5% 25 mM ammonium carbonate in ddH2O) as the aqueous phase and buffer B (99% ACN with 5% ddH2O) as the organic phase. The gradient ran from 99% buffer B to 85% over 2 min, to 75% over 3 min, to 60% over 3 min, then decreased to 40% and held for 1 min, then dropped to 1% and held for 1 min, and then returned to 99% for 0.1 min and was re-equilibrated for 5 min between runs. Data were acquired using high-resolution multiple reaction monitoring (MRM HR).

### Determining metabolite identity and data pretreatment

Mass spectrometry data were acquired by The Utah Metabolomics Core Facility using SCIEX OS Software and chromatogram integration was performed with SCIEX Analytics. Processed data were exported to Excel (Microsoft, Redmond WA), and subjected to statistical analysis via the Excel Data Analysis add-in. Metabolite lists were curated to include only analytes of interest. Identity assignments were based on comparison to an in-house library of pure standards and the Human Metabolome Database (hmdb.ca) (Wishart et al., 2022). Compound annotations relied on MS/MS spectra acquired by information-dependent acquisition (IDA) and SWATH, together with the inhouse standard library. Pretreatment of the data was as follows: missing values were imputed at one-fifth of the lowest detected signal for each metabolite; signal intensities were normalized using a sample-specific factor calculated as the ratio of each sample’s D_9_-carnitine intensity to the mean D_9_-carnitine intensity across all samples and QC replicates; metabolites exhibiting a coefficient of variation (CV) ≥ 30 % across QC samples were excluded from further analysis.

### Statistical analysis of metabolomics data

We performed all statistical analyses in R (v4.5.0; (R Core Team, 2025), using the packages of the tidyverse (v2.0.0; (Wickham et al., 2019) and drc (v3.0-1; (Ritz et al., 2015). We downloaded bioclimatic variables at 10-minute resolution from WorldClim 2 (Fick and Hijmans, 2017). To normalize metabolite data, we first applied a log_10_ transformation on the normalized intensity value of each metabolite, and then applied pareto scaling by subtracting the mean log_10_ intensity for each metabolite and dividing by the square root of its standard deviation (van den Berg et al., 2006). To visualize major shifts in the captured metabolome across the regions and treatments, we performed a principal component analysis on the pareto scaled data. To test whether variation in the metabolome predicts heat tolerance, we analyzed the association between PC scores and genotype-specific LT_50_. For each temperature separately (25°C control and 32°C heat-shocked samples), we fit linear models regressing the score of each principal component on the genotype-specific LT_50_; p-values were corrected for multiple comparisons using the Benjamini-Hochberg procedure. To assess the metabolic relatedness among samples, we performed hierarchical clustering on the pareto-scaled data using Pearson correlation-based distance. We then visualized coordinated changes across the 36 metabolites in a heatmap, clustering the metabolites in the same way. Dendrograms are shown for both axes, with sample annotations indicating region and temperature treatment.

### Differentially abundant metabolites

We identified differentially abundant metabolites by fitting a linear mixed-effects model to the normalized intensities of each metabolite (including the three redox couple ratios), with region, temperature, and their interaction as fixed effects and genotype as a random effect to account for baseline differences among genotypes. We adjusted p-values using the Benjamini-Hochberg procedure to account for multiple comparisons. Additionally, to isolate the effect of heat shock within region, for metabolites with a significant interaction, we preformed post hoc simple-effects contrasts of temperature within each region and adjusted for multiple comparisons with the Benjamini-Hochberg procedure.

### Genotype-level redox shift

For each genotype and redox couple (NADH/NAD^+^, NADPH/NADP^+^ and GSH/GSSG), we quantified the heat-induced shift in redox state as the difference between the pareto-scaled mean ratio at 32°C and 25°C. We estimated the error using a 95% confidence interval, which was calculated as 1.96 × standard error.

## Data availability

All metabolomics data and code used for statistical analysis are available on GitHub (https://github.com/tsoleary/redox_balance).

## Acknowledgements

We would like to thank Alexandra Lewis for her work phenotyping heat tolerance in embryos. Metabolomics analysis was performed at the Metabolomics Core Facility at the University of Utah. Mass spectrometry equipment was obtained through NCRR Shared Instrumentation Grant 1S10OD016232-01, 1S10OD018210-01A1 and 1S10OD021505-01. This work was funded by NSF grant IOS-1750322 to BLL.

